# Effective population size of X chromosomes and haplodiploids under cyclical parthenogenesis

**DOI:** 10.1101/2024.04.05.586733

**Authors:** Thomas J. Hitchcock

## Abstract

Many organisms can reproduce both sexually and asexually. Many such groups also exhibit ploidy differences between males and females during the sexual phase of their life, including aphids, nematodes, wasps, and rotifers. The constraints of combining asexual reproduction with asymmetric ploidy often results in X chromosomes (or haplodiploids) which show distinct transmission genetics from conventional systems. This may be expected to impact the effective population size; however, we currently lack analytical expressions for such life cycles. To remedy this, here we generate expressions for the effective population size of X chromosomes and haplodiploids under cyclical parthenogenesis. By first analysing a purely sexual scenario, we show how these distinct transmission genetics generate an effective population size substantially smaller than that of the autosomes, and how it may be further altered by aspects of sex-specific ecology, inbreeding, and non-random X elimination. Considering this inheritance system in an explicit population structure also demonstrates how genetic differentiation builds up differently between autosomes and X chromosomes. By introducing asexual reproduction into the model, we can see how low frequencies of sex cause these differences between autosomes and X chromosomes to dissipate. We discuss the relevance of these results to some different groups and consider future avenues for both empirical and theoretical work.

## Introduction

Many organisms reproduce both asexually and sexually (Bell 1982; Hebert 1987; Normark 2003; De Meeûs et al. 2007; Orive and Krueger-Hadfield 2021; Ross et al. 2022). Many of these organisms also experience ploidy differences between males and females during the sexual phase of their life cycle, whether this be certain chromosomes (aphids, gall midges, *Stronglyoides* nematodes), portions of chromosomes (*Strongyloides papillosus*), or even the whole genome (rotifers, cynipid wasps) (Table 1). An increasing body of work has explored how facultative sex will shape patterns of genetic diversity, and how we may be able use genetic data to make inferences about the natural history and ecology of these organisms, and make predictions about their evolutionary trajectories (Orive 1993; Balloux et al. 2003; Ceplitis 2003; Prugnolle, Liu, et al. 2005; Hartfield et al. 2016; Rouger et al. 2016; Hartfield 2021; Stoeckel et al. 2021). This can be of practical relevance as these groups contain neglected tropical diseases, veterinary pathogens, agricultural pests, and disease vectors (Blackman and Eastop 2000; De Meeûs et al. 2009; Thamsborg et al. 2017; Buonfrate et al. 2020).

**Table 1:** Summary of the major groups combining cyclical parthenogenesis and asymmetric ploidy, either in the form of sex chromosomes or haplodiploidy.

|  |  |
| --- | --- |
| XX/X0 | **Aphidomorpha:** Aphids, phylloxerans, and adelgids share a core common life cycle with a variable number of apomictic reproductive cycles through most of the year, followed by the production of sexual males and females in autumn, who produce overwintering eggs (Simon et al. 2002; Havill and Foottit 2007; Forneck and Huber 2009). Males in these groups are XO and females XX, the consequence of random X elimination during the production of male embryos (Blackman 1987; Manicardi et al. 2015; Li et al. 2023). Males undergo a non-standard and highly variable meiosis, in which nullo-X sperm are inviable, such that all sperm contain an X (Manicardi et al. 2015). |
|  | **Nematoda:** In the genus *Strongyloides*, individuals alternate between a free-living and parasitic stage. Parasitic females reproduce asexually, producing eggs which pass out of the host in faeces. Sex determination varies but is generally some form of genome elimination/diminution of the X chromosome in the males such that males are X0 and females XX (Streit 2017). Females either develop as sexual females (heterogonic cycle), or directly develop as new parasitic females (homogonic cycle). Sexual females mate with males, and all of these offspring are parasitic females (Viney 2017). Across these cases, it appears that all offspring produced sexually are female due to the programmed cell death of the nullo-X sperm (Dulovic et al. 2022). |
| Haplodiploidy | **Rotifera:** Monogonant rotifers contain periods of both asexual and sexual reproduction during their life. Within the life cycle there are amictic (asexual) females who produce diploid eggs parthenogenetically. These eggs will typically be new amictic females, however environmental conditions – such as population density - induce the production of mictic (sexual) females. Mictic females produce haploid eggs which if unfertilized become males and if fertilized give rise to resting eggs, which will then hatch into new amictic females (Gilbert 2020). |
|  | **Hymenoptera:** Two groups within the Cynipidae, the Cynipini and Pediaspidini, exhibit a bivoltine life cycle alternating between an asexual and sexual form. There are two different types of asexual female, gynephores and androphores. Gynephores produce diploid daughters parthenogenetically. Androphores produce haploid eggs meiotically, which will then develop into haploid sons. Sexual females produce haploid eggs which must be fertilized, giving rise to the new generation of asexual females. Although amongst these groups there are also species which deviate from this life cycle (Stone et al. 2002). |

However, less attention has been paid to the intersection between asexuality and sex chromosomes (or haplodiploidy). This is important because (1) many such facultatively sexual organisms exhibit such inheritance systems, and (2) often the constraints of asexuality result in these behaving differently to their obligately sexual counterparts (Hebert 1987; Normark 2003; Table 1). For example, in cyclically parthenogenetic aphids, sexual males and females are asexual clones of their mother, but in the production of male embryos one X chromosome is eliminated in the polar body, rendering them X0 (Blackman 1987; Manicardi et al. 2015; Figure 1). Similar transmission genetics, albeit mechanistically different are shared amongst groups of nematodes, wasps, and rotifers (Table 1). These non-standard transmission genetics result in an X chromosome which spends an equal fraction of its evolutionary time in sexual females and sexual males (in contrast to the 2:1 ratio of standard XX/X0 systems), with consequences for mutation rate, recombination rate, and effective population size (Jaquiéry et al. 2012; Jaquiéry et al. 2013; Klein et al. 2021). On this latter point, simulations tailored to the life cycle of pea aphids have suggested that the effective population size of the X and the autosomes should be similar (Jaquiéry et al. 2012). Yet we currently lack analytical expressions for these inheritance systems, and thus it is unclear how these results might extend outwith the specific scenarios simulated, and across the varied life cycles and ecologies observed in these groups.

**Figure 1:**
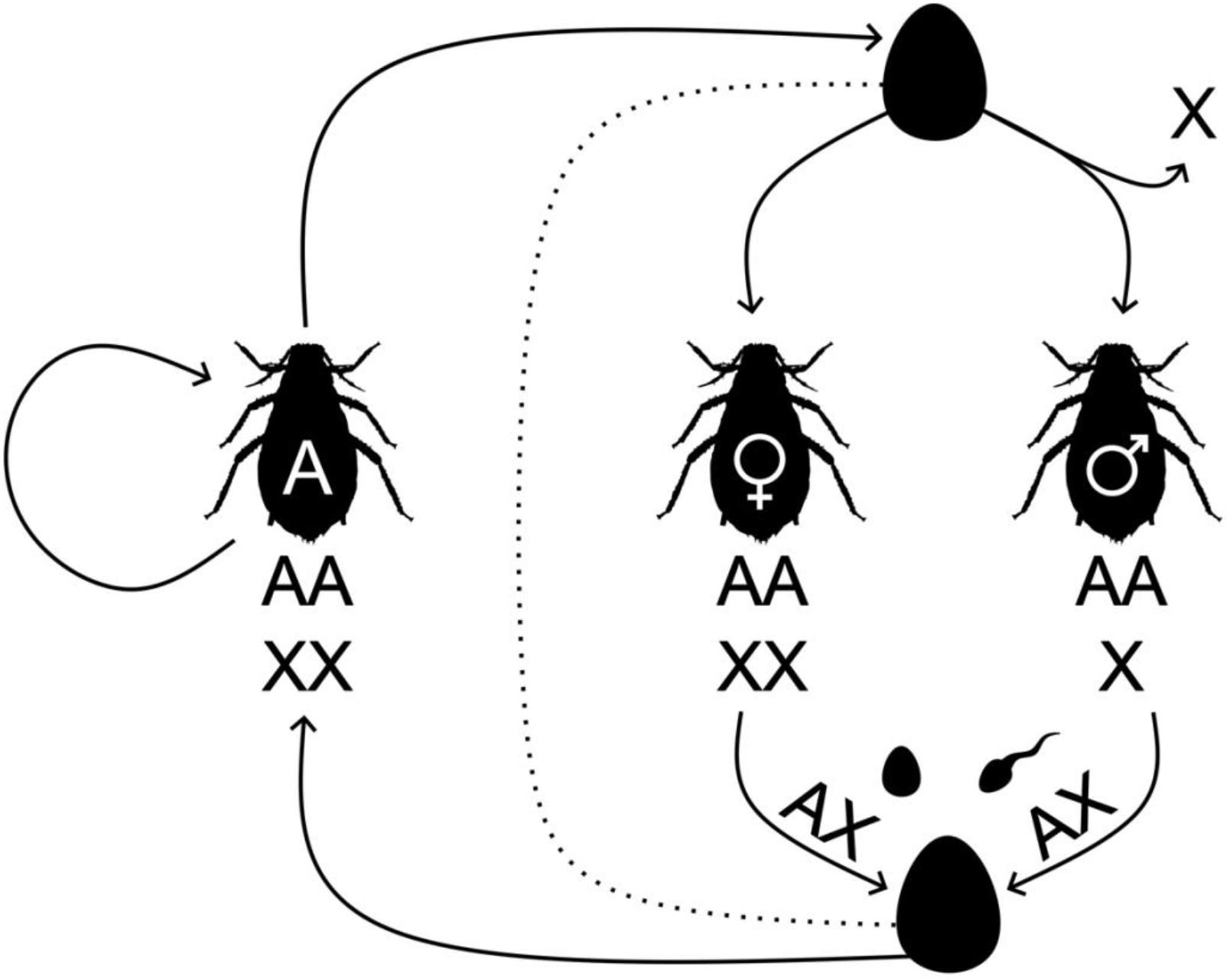
Simplified aphid life cycle. Random elimination of one of the X chromosomes determines the sex of an individual, producing XX females (♀) and X0 males (♂). Males only produce X containing sperm, and all offspring develop into XX asexual females (A) who clonally propagate for multiple generations before asexually producing sexual males and females once more. Here we also analyse a “pure sex” variant of this life cycle where sexually produced embryos skip the asexual phase entirely (dotted line).

To remedy this, here we generate expressions for the effective population size of X chromosomes and haplodiploids under cyclical parthenogenesis. We first analyse a purely sexual scenario, showing how the process of random X elimination results in an effective population size substantially smaller than that of the autosomes, and how it may be further altered by aspects of sex-specific ecology, inbreeding, and non-random X elimination. Next, considering this inheritance system in an explicit population structure demonstrates how *F*_*IS*_ and *F*_*ST*_ build up differently on autosomes and X chromosomes. Finally, introducing asexual reproduction into the model shows that with increasing amounts of asexuality results for X chromosomes and autosomes converge. We discuss the relevance of these results to some different groups and consider future avenues for both empirical and theoretical work.

### X elimination and effective population size

To fasten attention on this sex determination system and its effective population size consequences, we focus first on an obligately sexual scenario. All embryos are initially XX but those destined to become males eliminate one of their X chromosomes, such that they become X0 (Figure 1). Thus, we temporarily ignore the asexual phases that characterize these various groups (a scenario previously modelled in Klein et al. (2021)). We then compare the effective population size under random X elimination to a standard autosome and X chromosome. Provided that the population size is sufficiently large, such that gene lineages move between sexes at a much faster rate than coalescence, then we can calculate the effective population size by first calculating the average rate of coalescence across our various classes 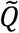, where the appropriate weights are class reproductive values, and then express the effective population size as 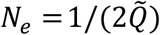 (Nordborg 1997; Laporte and Charlesworth 2002; Nordborg and Krone 2002; SM§1.1).

In the autosomal case, the probability that we sample two genes of maternal-origin is (1/2)^2^. If so, the probability they come from the same mother is 𝒜, and then they come from the same gene copy with probability (1/2). An identical logic applies to the paternal-origin route, but here we notate the probability of coming from the same father as ℬ. Putting this together, then our rate of coalescence for an autosomal locus will be:

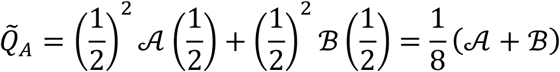

Assuming an even sex ratio and a Poisson distributed number of offspring (𝒜 = ℬ = 2/*N*), then our effective population size for an autosome will simply be 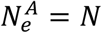.

Under random X elimination the logic is similar, with half of the gene copies of maternal-origin and half of paternal-origin. Now, however, when sampling from males there is only a single X chromosome that we can sample from, increasing the probability of coalescence:

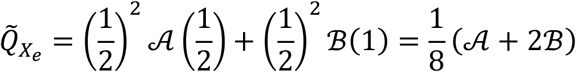

With the same assumptions as above, the effective population size becomes 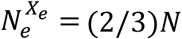. And for completeness, the rate of coalescence for a standard X chromosome system, whereby two thirds of copies are of maternal-origin, will be:

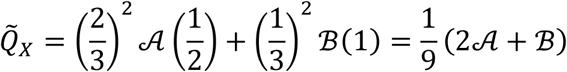

With the resultant effective population size being 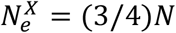. Thus, we can see that, perhaps contrary to expectations, a system whereby haploid males are generated through a process of random X elimination gives this portion of the genome an effective population size that is neither equivalent to a standard X, nor an autosomal system, but less than both.

### Sex-biased variance in reproductive success, inbreeding, and non-random X elimination

Other factors may alter these ratios of effective population size. For example, it is well understood that aspects of sex-specific demography can alter the 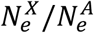ratio from the usual value of ¾ (Wright 1931; Wright 1933; Caballero 1995; Laporte and Charlesworth 2002). An increase in the probability of coming from the same mother 𝒜 (due to a higher variance in female reproductive success or fewer mothers) will decrease the relative effective population size of the X chromosome, whilst an increase in the probability of coming from the same father ℬ will tend to increase it (Figure 2). In contrast, from Equation 2, we can see that under random X elimination the opposite pattern will hold. Greater weight is now placed upon the probability of paternal sibship ℬ. Higher male variance in reproductive success (or male biased adult sex ratio) will reduce the relative effective population size, whilst higher female variance in reproductive success (or female biased adult sex ratio), will increase it (Figure 2).

**Figure 2:**
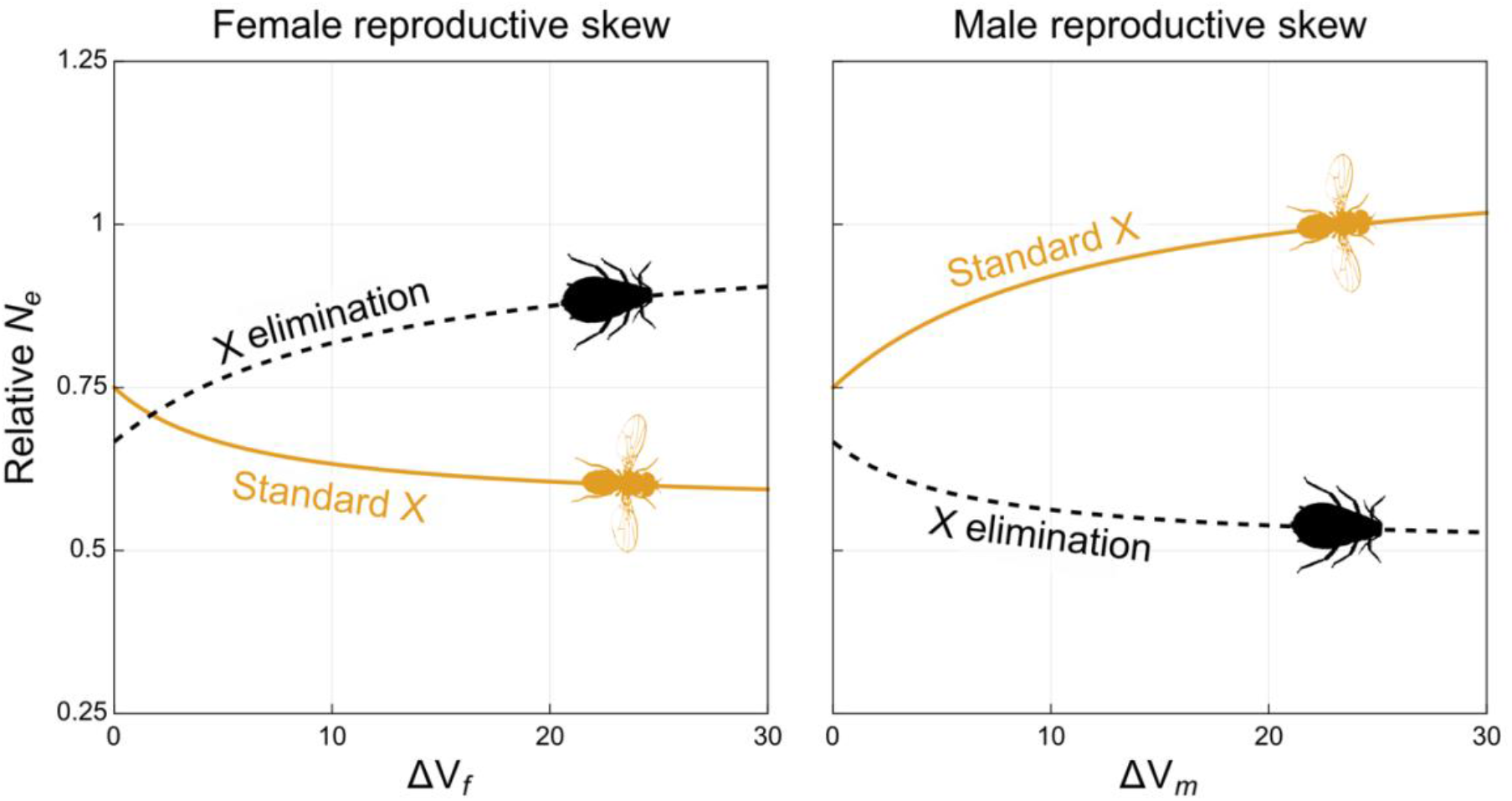
Relative effective population size of the X chromosome (*N*_*e,X*_/*N*_*e,A*_) under a standard X chromosome system, or random X elimination, when there is either female reproductive, or male reproductive skew. Δ*V*_f_ is the excess over the Poisson variance in offspring number for females, and Δ*V*_*m*_ is the equivalent value for males (Laporte & Charlesworth 2002, SM§1). In both cases an even sex ratio of breeders is assumed.

Inbredness will also impact the effective population size of sex chromosomes and autosomes differently (Caballero and Hill 1992; Wang 1996; Hedrick and Parker 1997; Laporte and Charlesworth 2002). If we denote the equilibrium inbreeding coefficient to be *F*, then for a diploid individual the probability of coalescence (given two gene copies descend from that individual) will be (1 + *F*)/2. For an autosomal locus, inbreeding will impact the rate of coalescence through both the maternal and paternal route. For X chromosome systems – due to the haploidy of males – inbreeding will only impact the maternal route. Additionally, as the maternal-origin route constitutes a greater fraction of the ancestry for a standard X chromosome, then inbreeding will have a more substantial impact here (and reduce the effective size more) than in a random X elimination system. Under random X elimination, with high levels of inbreeding (*F* → 1), the X and autosomes converge.

In some aphid species there is also evidence that the process of X elimination is non-random (Frantz et al. 2005; Monti et al. 2011; Wilson et al. 2014). Whilst on a population level there is no systematic bias towards maternal vs paternal elimination, individual females produce sons who are more likely to share the same X chromosome than expected by chance. Why this is occurs is not yet clear (Wilson et al. 2014). To model this effect, I assign a fraction *ϕ* of females whose offspring will undergo non-random elimination. In half of these individuals, their sons will always eliminate the maternal-origin X, and in the other half their sons will always eliminate the paternal-origin X (SM§1.2). If the mating system is monogamous, then the effective population size of the X becomes:

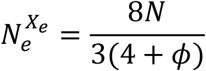

As the degree of non-random elimination increases, the effective population size falls. As *ϕ* → 1, the effective population size of the X chromosome will be just over half that of the autosomes. With increasing amounts of polyandry however, the magnitude of this effect is reduced (SM§1.2).

### Population structure

Next, we turn our attention to the consequences of population structure. In previous analyses it has been shown that, when there is spatial structure within populations, genetic differentiation may build up differently on autosomes and sex chromosomes (Laporte and Charlesworth 2002). In such cases, *F*_*ST*_ typically builds up more rapidly on the X chromosomes than the autosomes. To explore this phenomenon with X elimination, we analyse an infinite island model, in which the various patches are connected by dispersal (Wright 1931; Rousset 2004; Prugnolle et al. 2005b). The life cycle begins with the production of sexual juveniles on the patch. Those juveniles then disperse with sex-specific probabilities to new patches, whereupon there is density dependent competition, those individuals develop into adults, and then the life cycle begins once more.

In the infinite island model (and under the infinite allele model), *F*_*ST*_ is equivalent to the consanguinity between two gene copies sampled in different individuals on the same patch *ρ*_1_, as the consanguinity between two individuals on separate patches *ρ*_2_ is assumed to be zero:

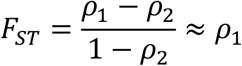

By solving the recursion equations for our consanguinities at equilibrium we find that the *F*_*ST*_ between juveniles (pre-dispersal) on a patch in the autosomal case is given by:

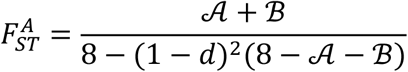

Whilst under random X elimination our expression is instead:

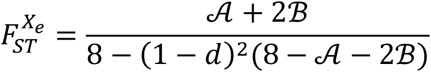

As 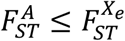, then for all dispersal parameters and patterns of reproductive skew, we expect more pronounced genetic differentiation on an aphid-like X than on the autosomes (Figure 3). Similarly, *F*_*IS*_ takes on a larger (more negative) value under random X elimination than on the autosomes. Moreover, unlike on standard X chromosomes, whilst sex-biased dispersal per se does change values of *F*_*ST*_, the specific direction of sex bias (male vs female biased dispersal) has no impact (SM§2).

**Figure 3:**
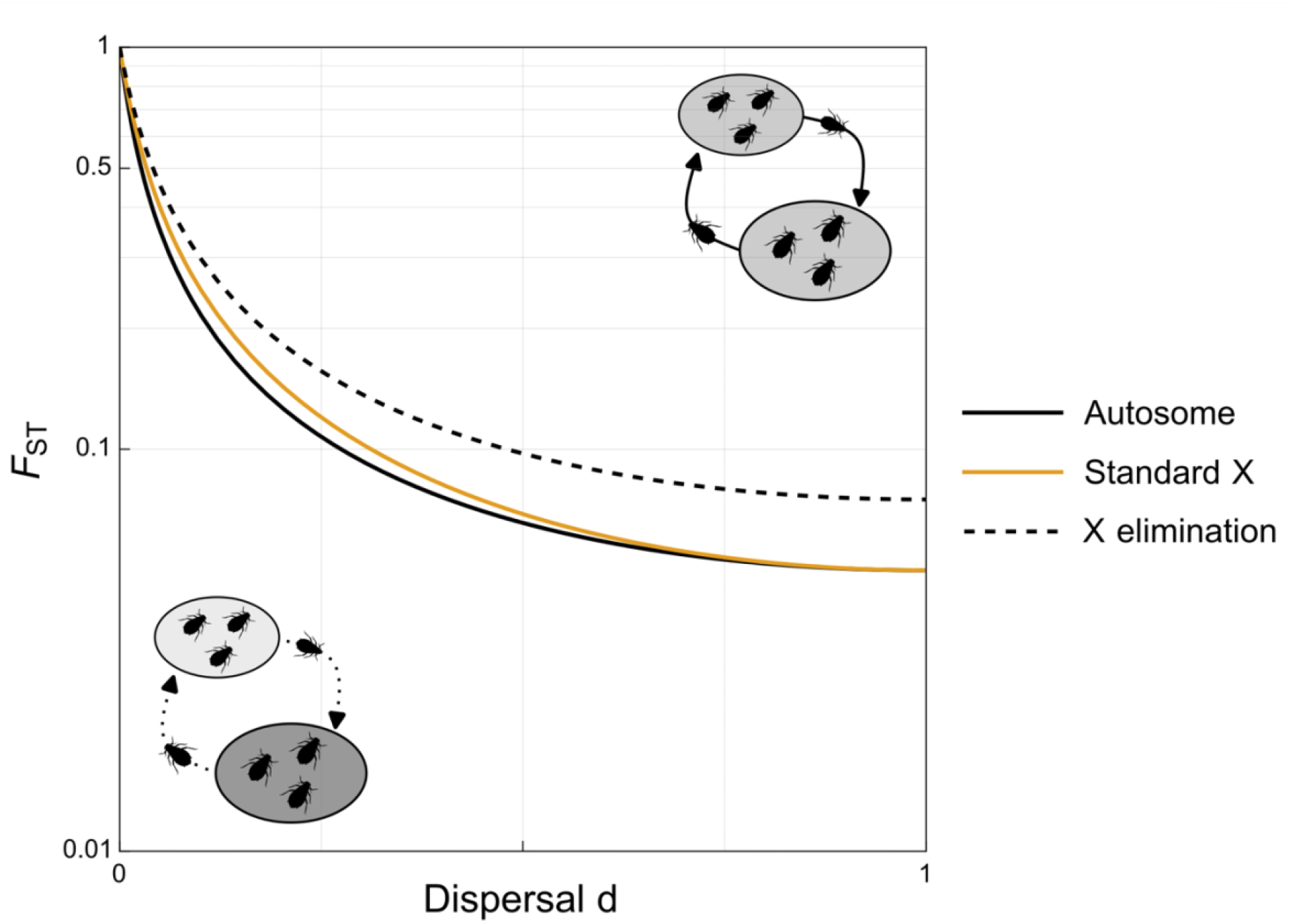
Population structure results in more substantial genetic differentiation under random X elimination than on autosomes. Here *F*_*ST*_ between male and female juveniles in an infinite island model is plotted as a function of dispersal rates. Dispersal rates are equal between males and females (*d*_*f*_ = *d*_*m*_ = *d*) and there are 10 males and females per patch (𝒜 = ℬ = 1/10). See SM§2 for further details.

### Asexual reproduction

Thus far, we have assumed obligate sexuality. However, all the groups which share this inheritance system contain asexual phases within their life cycle (Table 1.). To incorporate this, we now consider an asexually produced generation and ask what the probability of coalescence, 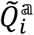, is. We label the two gene copies in an asexual female based on the sexual parent they originally descended from (either maternal-origin and paternal-origin). In our asexually produced offspring, we sample two maternal-origin descendant genes with probability (1/2)^2^. If so, then they descend from the same (asexual) mother with probability *𝒫*, in which case they came from the same gene copy.

Following the same procedure for the descendants of the paternal-origin copy and assembling give us:

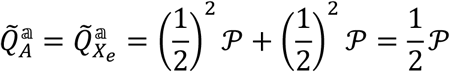

Which applies to both the autosomal case and the X elimination case. For a standard X chromosome system, the scenario is slightly different. Whilst physically ½ of the copies in the offspring are of paternal-origin, the eventual consequences of sex mean that these gene copies are only half as valuable as their maternal-origin counterparts, and so:

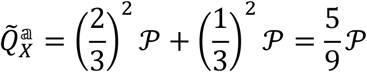

If *σ* is the frequency of sexual transitions, and we assume that sex occurs at a rate much higher than coalescence, then we can approximate the rate of coalescence as:

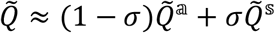

Where 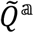 is our coalescent rate for asexual transitions, and 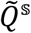 for sexual ones (Hartfield et al. 2016; SM§3).

We can see that as the autosomes and X elimination have an identical rate of coalescence during the asexual phase, then increasing fractions of asexual reproduction will make their effective population sizes more alike and as the rate of sex *σ* tends to zero, then they will converge exactly. Conversely, when there are high levels of sexual reproduction then they will be quite different. In a similar fashion, the different signatures of *F*_*ST*_ and *F*_*IS*_ on aphid-like X chromosomes also dissipate with increasing rates of asexuality (SM§2). In contrast, if we combined a standard X chromosome with asexuality (this could occur if the paternal-origin X systematically eliminated), then whilst asexuality would make the effective population size of the X and autosomes more similar, provided sex does still occur then coalescent rate will always be higher and effective population size lower (Figure 4).

**Figure 4:**
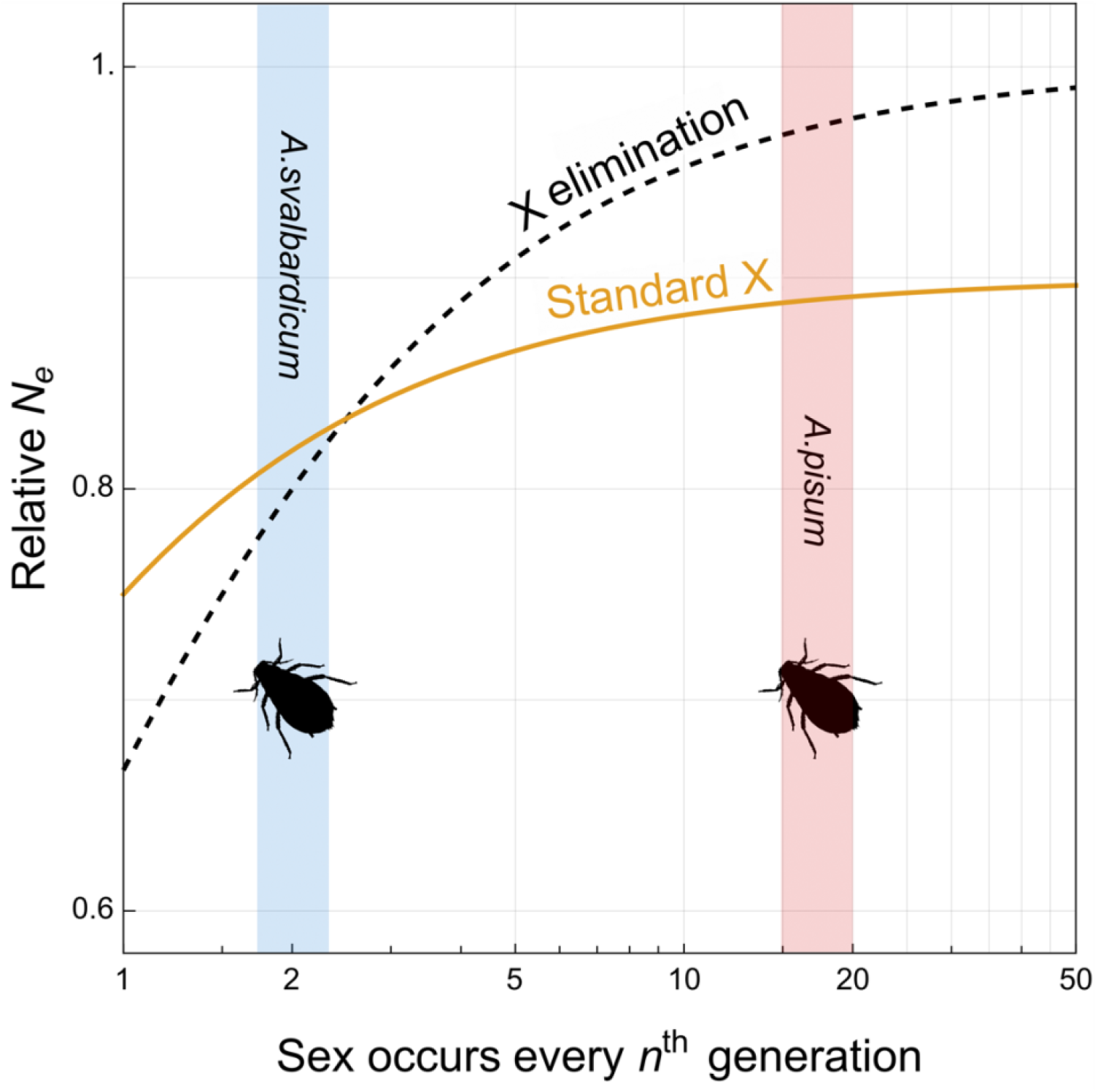
Relative effective population size of the X chromosome (*N*_*e,X*_/*N*_*e,A*_) under a standard X like inheritance system, or a random X elimination system, when there are differing numbers of asexual generations in the life cycle. Two aphid species – the pea aphid (*Acyrthosiphon pisum*) and high arctic aphid (*Acyrthosiphon svalbardicum*) - with differing frequencies of sexual vs asexual reproduction are highlighted (Strathdee et al. 1993; Jaquiéry et al. 2012).

## Discussion

Here we have shown that, despite the equivalent amounts of time X chromosomes and autosomes spend in males and females under cyclical parthenogenesis, X chromosomes may nonetheless experience a reduced effective population size, due the effects of sampling from haploid males. This extent of this will be altered by further aspects of the mating scheme and ecology, such as inbreeding, the sex ratio, and sex-biased variance in reproductive success. Most notably, increasing levels of asexuality will reduce differences between the autosomes and X chromosomes, such that with infrequent sex their effective population sizes will be very similar, which matches the pattern previously seen in simulations (Jaquiéry et al. 2012).

One signature of this effect may be seen in patterns of neutral genetic diversity. From these results we should always expect less diversity on the X chromosomes than the autosomes, and with the most pronounced differences in those populations with frequent sexual reproduction. Aphids may provide good systems to test this hypothesis, and there are an increasing number of aphid species (and close relatives) which now have chromosome level genome assemblies (Mathers et al. 2021; Dial et al. 2023; Li et al. 2023). Most aphids must reproduce sexually to produce overwintering eggs, but the number of possible asexual generations scales with the length of the breeding season (Simon et al. 2002). Consequently, the frequency of sexual vs asexual reproduction may well scale with latitude and/or altitude. The relative neutral genetic diversity on these two portions of the genome should also scale in a similar fashion, being more similar in more temperate climes (with longer asexual breeding seasons), and more divergent in areas nearer the poles or of high altitude (with shorter asexual breeding seasons). Although this pattern might be slightly complicated by the presence of (and gene flow from) obligately parthenogenetic lineages, which themselves are often concentrated in more temperate areas (Martel et al. 2020; Rimbault et al. 2023).

Differences in patterns of genetic diversity between the X and the autosomes might also be used to infer rates of sex, or sex specific demography in natural populations. This may be particularly valuable in species where these are hard to assay directly, and sex is relatively frequent. In *Strongyloides* nematodes, the relative frequency of sex is predominantly governed by the decision of females to either directly develop as infectious larvae (homogonic) or develop instead into free living, sexual females (heterogonic development) (Streit 2008). Experimental work has shown environmental factors can influence the fraction that pass through these two routes, such as temperature and immune response of the host (Harvey et al. 2000; Viney and Lok 2015). Moreover, rates of sex are thought to vary between species and strains of *Strongyloides* (Viney et al. 1992). Comparisons between the X and the autosomes in these systems could allow inferences for the frequency of these two different pathways in natural systems. Although in groups where rates of asexuality appear very high then this comparison will be less informative (Cole et al. 2023).

In addition to neutral processes, the differences in effective population size and ploidy may also impact rates of adaptation, altering the probabilities that beneficial (and deleterious) variants fix (Meisel and Connallon 2013). There is some evidence for a faster X effect in aphids, although this appears to be due primarily driven by the lower expression of genes on the X chromosome, and thus increased fixation probability of slightly deleterious alleles (Jaquiéry et al. 2018). In general, it is not clear how the balance between the haploid expression observed in the males and the lower effective population size will balance out with shifting rates of sex. Additionally, this will be sensitive to both patterns of sex-biased gene expression (which will only affect the marginal fitness effects), and sex-biased patterns of reproductive success (which will only affect the effective population size). An interesting possible comparison in this regard are the sex chromosomes of *Strongyloides papillosus*. Here, whilst an entire X chromosome is eventually eliminated, a portion is retained in the soma, a consequence of an ancestral X-A fusion event (Nemetschke et al. 2010). Thus, whilst the entirety of this chromosome shares the same effective population size, only a portion experiences the effects of haploid selection, making it a good candidate for comparative work.

Whilst we have focused on broad similarities across these life cycles, it is worth emphasizing the diversity in life cycle structures amongst these groups. For example, in the best studied of the cynipid wasps there are two distinct types of female – gynandromorphs and andromorphs – which exclusively produce either sons or daughters, a type of split sex ratio system, although distinct from that seen in other Hymenoptera or Diptera (Stone et al. 2002; Meunier et al. 2008; Baird et al. 2023). Differences in the proportions of these morphs, or differences in reproductive skew between them, will further reduce the effective population size. Similarly, in the monogononant rotifers males are not directly produced by the asexual (amictic) females, but instead produced by unfertilized (mictic females)(Gilbert 2020). This generates an extra cryptic generation through the paternal line in these groups, which may well result in a form of systemic, higher reproductive skew through the paternal line. Finally, whilst we have not explicitly discussed polyembryony here (as this is not typically treated under the umbrella of cyclical parthenogenesis), it shares many of the same life cycle features (Craig et al. 1997). Although it is important to note that haplodiploid polyembryonic species (such as certain parasitoid wasps) typically demonstrate “true” haplodiploid inheritance (Segoli et al. 2010), unlike the oak gall wasps and monogonont rotifers. Collectively, the core patterns we have outlined here combined with the rich diversity of ecologies and life cycles seen within these groups may make them valuable study systems for comparative genomics, and potentially rich sources of future empirical and theoretical investigation.

## Funding and Acknowledgements

This work was supported by the Special Postdoctoral Researchers Program funded by RIKEN. I thank R. Iritani, J. Jaquiéry, and L. Ross for helpful discussion and feedback.

