## Supplementary Material for "Effective population size of X chromosomes and haplodiploids under cyclical parthenogenesis"

Thomas J. Hitchcock[1*]

[1]RIKEN Interdisciplinary Theoretical and Mathematical Sciences Program (iTHEMS), Wako, Saitama, Japan

#### 1 Fast-time scale approximation of the coalescent

##### 1.1 Pure sexual reproduction

First, we consider the pure sexual scenario. Here we may write out a weighted probability of coalescence as $\tilde{Q}$, and the effective population size as $N_e = \frac{1}{2\tilde{Q}}$. $\tilde{Q}$ is calculated as:

$$\tilde{Q} = \sum_k \sum_i \sum_j c_{\mathrm{k}\to\mathrm{i}} c_{\mathrm{k}\to\mathrm{j}} Q^{\mathrm{k}}_{\mathrm{ij}} \tag{S1}$$

Where $c_{\mathrm{k}\to\mathrm{j}}$ is the reproductive value of the transition between states $k$ and $i$, and $Q^{\mathrm{k}}_{\mathrm{ij}}$ is the probability that two gene lineages in $i$ and $j$ coalesce given that they both came from state $k$.

The reproductive values of the transitions are calculated by first taking the left-eigenvalue of the transition matrix $T = (t_{\mathrm{ij}})$, where $t_{ij}$ is the probability that a gene copy in state $i$ came from state $j$ in the previous generation. If we let $\alpha$ be the probability that a gene in female came from a female in the previous generation, and $\beta$ be the probability that a gene in a male came from a male in the previous generation then:

$$T = \begin{pmatrix} \alpha & 1-\alpha \\ 1-\beta & \beta \end{pmatrix} \tag{S2}$$

And so our class reproductive values of males and females are:

$$\begin{pmatrix} c_{\mathrm{f}} & c_{\mathrm{m}} \end{pmatrix} = \begin{pmatrix} c_{\mathrm{f}} & c_{\mathrm{m}} \end{pmatrix} . T \tag{S3a}$$

$$\begin{pmatrix} c_{\mathrm{f}} & c_{\mathrm{m}} \end{pmatrix} = \begin{pmatrix} \frac{1-\beta}{2-\alpha-\beta} & \frac{1-\alpha}{2-\alpha-\beta} \end{pmatrix} \tag{S3b}$$

And then the values of the transitions are:

$$E = \begin{pmatrix} c_{\mathrm{f}\to\mathrm{f}} & c_{\mathrm{f}\to\mathrm{m}} \\ c_{\mathrm{m}\to\mathrm{f}} & c_{\mathrm{m}\to\mathrm{m}} \end{pmatrix} = \begin{pmatrix} \frac{\alpha(1-\beta)}{2-\alpha-\beta} & \frac{(1-\alpha)(1-\beta)}{2-\alpha-\beta} \\ \frac{(1-\alpha)(1-\beta)}{2-\alpha-\beta} & \frac{(1-\alpha)\beta}{2-\alpha-\beta} \end{pmatrix} \tag{S4}$$

For both a standard autosomal case $A$, and the random X-elimination case (which we will refer to as an "aphid-like" X chromosome) $X_a$, $\alpha = \beta = 1/2$. For a standard X chromosome system $X$, $\alpha = 1/2$ and $\beta = 0$.

Next, we calculate the probabilities that two genes copies are identical by descent, $Q^{\mathrm{k}}_{\mathrm{ij}}$. For a two sex system, there are 6 possible pairs of consanguinities. Following Laporte & Charlesworth (2002), we can write the various probabilities of consanguinity as:

$$Q^{\mathrm{f}}_{\mathrm{ff}} = \mathcal{A}_{\mathrm{ff}}\mu = \left(\frac{1 + \left(\Delta V^{\mathrm{f}}_{\mathrm{f}} / n^2_{\mathrm{ff}}\right)}{N_{\mathrm{f}}}\right)\mu \tag{S5a}$$

$$Q^{\mathrm{f}}_{\mathrm{fm}} = \mathcal{A}_{\mathrm{fm}}\mu = \left(\frac{1 + \left(\Delta C^{\mathrm{f}}_{\mathrm{fm}} / n_{\mathrm{ff}} n_{\mathrm{fm}}\right)}{N_{\mathrm{f}}}\right)\mu \tag{S5b}$$

$$Q^{\mathrm{f}}_{\mathrm{mm}} = \mathcal{A}_{\mathrm{mm}}\mu = \left(\frac{1 + \left(\Delta V^{\mathrm{f}}_{\mathrm{m}} / n^2_{\mathrm{fm}}\right)}{N_{\mathrm{f}}}\right)\mu \tag{S5c}$$

$$Q^{\mathrm{m}}_{\mathrm{ff}} = \mathcal{B}_{\mathrm{ff}}\nu = \left(\frac{1 + \left(\Delta V^{\mathrm{m}}_{\mathrm{f}} / n^2_{\mathrm{mf}}\right)}{N_{\mathrm{m}}}\right)\nu \tag{S5d}$$

$$Q^{\mathrm{m}}_{\mathrm{fm}} = \mathcal{B}_{\mathrm{fm}}\nu = \left(\frac{1 + \left(\Delta C^{\mathrm{m}}_{\mathrm{fm}} / n_{\mathrm{mf}} n_{\mathrm{mm}}\right)}{N_{\mathrm{m}}}\right)\nu \tag{S5e}$$

$$Q^{\mathrm{m}}_{\mathrm{mm}} = \mathcal{B}_{\mathrm{mm}}\nu = \left(\frac{1 + \left(\Delta V^{\mathrm{m}}_{\mathrm{m}} / n^2_{\mathrm{mm}}\right)}{N_{\mathrm{m}}}\right)\nu \tag{S5f}$$

Where $\mathcal{A}_{\mathrm{ij}}$ is the probability that gene copies from sexes $i$ and $j$ came from the same mother (given they are both of maternal-origin) and $\mathcal{B}_{\mathrm{ij}}$ is the probability that gene copies from sexes $i$ and $j$ came from the same father (given they are both of paternal-origin). The probability that they are identical by descent given this is then $\mu$ for offspring of females and $\nu$ for offspring from males.

We can rewrite this in terms of excesses from the Poisson variance in offspring number, where the deviation in the variance of offspring of sex $i$ from a parent of sex $j$ is $\Delta V^{\mathrm{j}}_{\mathrm{i}}$, and the mean number of offspring of sex $i$ from a parent of sex $j$ is $n_{\mathrm{ji}}$, and $N_{\mathrm{i}}$ is the number of parents of sex $i$ (Laporte and Charlesworth, 2002). Similarly, the deviation of the covariance in the production of sons and daughters from the Poisson covariance (of 0), is written as $\Delta C^{\mathrm{f}}_{\mathrm{fm}}$.

If there is a sex ratio of $c$ which is consistent across breeding individuals, then these expressions can be neatly reduced down to:

$$\tilde{Q} = c^2_{\mathrm{f}}\mathcal{A}\mu + c^2_{\mathrm{m}}\mathcal{B}\nu \tag{S6}$$

Where:

$$\mathcal{A} = \frac{1 + (1-c)^2 \Delta V_{\mathrm{f}}}{N_{\mathrm{f}}} \tag{S7a}$$

$$\mathcal{B} = \frac{1 + c^2 \Delta V_{\mathrm{m}}}{N_{\mathrm{m}}} \tag{S7b}$$

Where the class reproductive values, and probabilities of coalescence within individuals will depend on our genetic system of interest:

$$\tilde{Q}_A = \left(\frac{1}{2}\right)^2 \mathcal{A}\left(\frac{1+\mathcal{F}}{2}\right) + \left(\frac{1}{2}\right)^2 \mathcal{B}\left(\frac{1+\mathcal{F}}{2}\right) = \frac{1}{8}(1+\mathcal{F})(\mathcal{A}+\mathcal{B}) \tag{S8a}$$

$$\tilde{Q}_{X_a} = \left(\frac{1}{2}\right)^2 \mathcal{A}\left(\frac{1+\mathcal{F}}{2}\right) + \left(\frac{1}{2}\right)^2 \mathcal{B} = \frac{1}{8}(\mathcal{A}(1+\mathcal{F}) + 2\mathcal{B}) \tag{S8b}$$

$$\tilde{Q}_{X} = \left(\frac{2}{3}\right)^2 \mathcal{A}\left(\frac{1+\mathcal{F}}{2}\right) + \left(\frac{1}{3}\right)^2 \mathcal{B} = \frac{1}{9}(2\mathcal{A}(1+\mathcal{F}) + \mathcal{B}) \tag{S8c}$$

We can see the asymmetric effect that inbreeding, sex ratio, and sex-biased variance in reproductive success will have upon the rate of coalescence, and thus effective population size.

#### 1.2 Non-random X elimination

One complication to the above scenario occurs when there is non-random elimination of the X chromosome in males. To incorporate this - but retain an equal distribution of maternal and paternal elimination - we assign a fraction of females $\phi$ to have non-random elimination, of which half have sons that exclusively eliminate their maternal-origin X chromosome, and the other half eliminate their paternal-origin X chromosome. For simplicity, we assume a Poisson variance in offspring number, and an equal sex ratio.

Given we sample two males whose X chromosomes are of maternal-origin, the probability they come from the same mother is:

$$\mathcal{A}_{\mathrm{mm}} = \phi^2 \frac{2}{\phi N_{\mathrm{f}}} + (1-\phi)^2 \frac{1}{(1-\phi)N_{\mathrm{f}}} = \frac{1+\phi}{N_{\mathrm{f}}} \tag{S9}$$

Depending on the mating scheme, the probability of consanguinity through fathers may also be increased. If mating is monogamous then $\mathscr{B}_{\mathrm{mm}} = \left(1+\phi\right)/N_{\mathrm{m}}$, whilst if every male mated with every female then the result would be $\mathscr{B}_{\mathrm{mm}} = 1/N_{\mathrm{m}}$. Let us scale between these two scenarios with $\lambda$ such that:

$$\mathcal{B}_{\mathrm{mm}} = (1-\lambda)\frac{1}{N_{\mathrm{m}}} + \lambda \frac{1+\phi}{N_{\mathrm{m}}} = \frac{1+\lambda\phi}{N_{\mathrm{m}}} \tag{S10}$$

Inserting these into the equations from before, gives us:

$$\tilde{Q}_{X_a} = \frac{12+\phi(1+2\lambda)}{16N} \tag{S11}$$

And so:

$$N_e^{X_a} = \frac{8N}{12+\phi(1+2\lambda)} \tag{S12}$$

If $\phi = 1$ and $\lambda = 1$ then this reduces the effective population size by $1/5$ of its original value.

#### 1.3 Mixed sexual and asexual reproduction

Now let us introduce asexuality into the model. Let a proportion $1-\sigma$ of generations be asexual and the probability of coalescence given you come through an asexual transition be $\tilde{Q}^{\mathbb{a}}$. Similarly, let a fraction $\sigma$ of generations be sexual, and during a sexual transition the probability of coalescence be $\tilde{Q}^{\mathbb{s}}$. One can then approximate the rate of coalescence as:

$$\tilde{Q} \approx (1-\sigma)\tilde{Q}^{\mathbb{a}} + \sigma\tilde{Q}^{\mathbb{s}} \tag{S13}$$

We have already written our the coalescence probabilities for our sexual transitions above. For the asexual transitions, label the gene copies in our current generation by whether, during the last sexual transition, they came through the maternal route $c_{\mathbb{m}}$ or the paternal route $c_{\mathbb{p}}$. We will assume that asexual reproduction is clonal, such that gene copies cannot effectively shuffle between these states during the asexual phase. In order for two genes then to coalescence, both must have come through the same route, either both maternal $c_{\mathbb{m}}^2$ or both paternal $c_{\mathbb{p}}^2$. If two gene copies come from the same route, then the probability that they descend from the same asexual mother is $\mathcal{P}$. Putting this together:

$$\tilde{Q}^{\mathbb{a}} = c_{\mathbb{m}}^2 \mathcal{P} + c_{\mathbb{p}}^2 \mathcal{P} \tag{S14}$$

For the autosomes and aphid-like X chromosomes $c_{\mathbb{m}} = c_{\mathbb{p}} = 1/2$. In contrast, for our standard X chromosome, due to the fact that paternal-origin X chromosomes will eventually be disposed of in males $c_{\mathbb{m}} = 2c_{\mathbb{p}} = 2/3$.

#### 2 F-statistics in the infinite island model

Here we model the same inheritance system within the context of an infinite island model, and from this calculate the values of $F_{\mathrm{IS}}$ and $F_{\mathrm{ST}}$. These F-statistics can can be expressed in terms of consanguinities (Cockerham, 1969; Cockerham, 1973; Rousset, 2004):

$$F_{\mathrm{IS}} = \frac{\rho_0 - \rho_1}{1 - \rho_1} \tag{S15a}$$

$$F_{\mathrm{ST}} = \frac{\rho_1 - \rho_2}{1 - \rho_2} \tag{S15b}$$

Moreover, these values may differ based on whether males or females are sampled, particularly, if there is sex-specific migration patterns, reproductive skew, or asymmetric inheritance systems (Vitalis, 2002).

We analyse a life-cycle similar to that of Prugnolle et al. (2005), in which we begin with a population of adults, who first may disperse between patches with sex-specific probabilities. They then mate, their offspring then clonally reproduce, and they give rise to the new generation of adult sexuals. An important difference between our implementation here and the life cycle analysed by Prugnolle and colleagues is that, unlike in trematodes, the groups of interest here have an asexual form that can give rise to both male and female sexuals.

First, we can write out the vector:

$$\boldsymbol{\rho}_{\mathrm{j}} = \begin{pmatrix} \rho_0 \\ \rho_1^{\mathrm{ff}} \\ \rho_1^{\mathrm{fm}} \\ \rho_1^{\mathrm{mm}} \end{pmatrix} \tag{S16}$$

Which denote the consanguinity between two gene copies within an individual ($\rho_0$), or two gene copies in different individuals ($\rho_1$), with the censusing occurring at the birth of new sexual juveniles. Those juveniles then may disperse with sex-specific probabilities between patches before maturing into adults, such that the consanguinity between adults $\boldsymbol{\rho}_{\mathrm{a}}$ is:

$$\boldsymbol{\rho}_{\mathrm{a}} = \boldsymbol{M}.\boldsymbol{\rho}_{\mathrm{j}} \tag{S17}$$

Where $\boldsymbol{M}$ is the migration matrix, where $h_{\mathrm{f}}$ and $h_{\mathrm{m}}$ are the probabilities that females and males respectively remain on the natal patch.

$$\boldsymbol{M} = \begin{pmatrix} 1 & 0 & 0 & 0 \\ 0 & h_\mathrm{f}^2 & 0 & 0 \\ 0 & 0 & h_\mathrm{f} h_\mathrm{m} & 0 \\ 0 & 0 & 0 & h_\mathrm{m}^2 \end{pmatrix} \tag{S18}$$

Adults then mate, producing the new "asexual" clonal lineages. The consanguinities between these clonal individuals $\boldsymbol{\rho}_\mathrm{c}$ is thus:

$$\boldsymbol{\rho}_\mathrm{c} = \boldsymbol{S}.\boldsymbol{\rho}_\mathrm{a} + \boldsymbol{S}_\mathrm{c} \tag{S19}$$

Where:

$$\boldsymbol{S} = \begin{pmatrix} 0 & 0 & 1 & 0 \\ \alpha^2 \mathcal{A}_\mathrm{ff}(1-\mu) + (1-\alpha)^2 \mathcal{B}_\mathrm{ff}(1-\nu) & \alpha^2 (1-\mathcal{A}_\mathrm{ff}) & 2\alpha(1-\alpha) & (1-\alpha)^2(1-\mathcal{B}_\mathrm{ff}) \\ \alpha(1-\beta)\mathcal{A}_\mathrm{fm}(1-\mu) + (1-\alpha)\beta\mathcal{B}_\mathrm{fm}(1-\nu) & \alpha(1-\beta)(1-\mathcal{A}_\mathrm{fm}) & \alpha\beta + (1-\alpha)(1-\beta) & (1-\alpha)\beta(1-\mathcal{B}_\mathrm{fm}) \\ (1-\beta)^2 \mathcal{A}_\mathrm{mm}(1-\mu) + \beta^2 \mathcal{B}_\mathrm{mm}(1-\nu) & (1-\beta)^2(1-\mathcal{A}_\mathrm{mm}) & 2(1-\beta)\beta & \beta^2(1-\mathcal{B}_\mathrm{mm}) \end{pmatrix} \tag{S20}$$

And:

$$\boldsymbol{S}_\mathrm{c} = \begin{pmatrix} 0 \\ \alpha^2 \mathcal{A}_\mathrm{ff}\mu + (1-\alpha)^2 \mathcal{B}_\mathrm{ff}\nu \\ \alpha(1-\beta)\mathcal{A}_\mathrm{fm}\mu + (1-\alpha)\beta\mathcal{B}_\mathrm{fm}\nu \\ \beta^2 \mathcal{A}_\mathrm{mm}\nu + (1-\beta)^2 \mathcal{B}_\mathrm{mm}\mu \end{pmatrix} \tag{S21}$$

Where $\alpha$ is the probability that a gene in a female descends from a female, and $\beta$ is the probability a gene copy in a male descends from a male. The probability of an individual of sex $i$ and $j$ descending from the same mother is $\mathcal{A}_\mathrm{ij}$, and from the same father as $\mathcal{B}_\mathrm{ij}$, given both copies are of maternal or paternal-origin respectively. $\mu$ and $\nu$ are defined slightly differently to above, instead as the probabilities of coming from the same physical copy in the mother or father respectively, given one comes from the same parent.Those individuals then clonal replicate, before producing the new sexual juveniles:

$$\boldsymbol{\rho}_\mathrm{j}' = \boldsymbol{A}.\boldsymbol{\rho}_\mathrm{c} + \boldsymbol{A}_\mathrm{c} \tag{S22}$$

Where:

$$\boldsymbol{A} = \begin{pmatrix} 1 & 0 & 0 & 0 \\ 2\alpha(1-\alpha)\mathcal{P} & (1-\mathcal{P}) & 0 & 0 \\ (\alpha\beta + (1-\alpha)(1-\beta))\mathcal{P} & 0 & (1-\mathcal{P}) & 0 \\ 2(1-\beta)\beta\mathcal{P} & 0 & 0 & (1-\mathcal{P}) \end{pmatrix} \tag{S23}$$

And:

$$\boldsymbol{A}_\mathrm{c} = \begin{pmatrix} 0 \\ \left(\alpha^2 + (1-\alpha)^2\right)\mathcal{P} \\ \mathcal{P}(\alpha(1-\beta) + (1-\alpha)\beta) \\ \left(\beta^2 + (1-\beta)^2\right)\mathcal{P} \end{pmatrix} \tag{S24}$$

Where $\mathcal{P}$ is defined as the probability that two juveniles descend from the same clonal lineage. Thus there may be many rounds of clonal reproduction during this phase, with $\mathcal{P}$ describing the eventual increase in the probability of sibship arising from this reproduction. Alternatively, one can eliminate the effects of clonal reproduction entirely by setting $\mathcal{P} = 0$, which simply returns us to the pure sexual scenario. We can then solve for the quasi-equilibrium value of $\hat{\boldsymbol{\rho}}_{\mathrm{j}}$.

$$\hat{\boldsymbol{\rho}}_{\mathrm{j}} = \boldsymbol{A}.\boldsymbol{S}.\boldsymbol{M}.\hat{\boldsymbol{\rho}}_{\mathrm{j}} + \boldsymbol{A}.\boldsymbol{S}_c + \boldsymbol{A}_c \tag{S25}$$

In general, full expressions for the consanguinities are a little cumbersome, however, we can highlight some specific results. First, when $\mathcal{A} = \mathcal{A}_{\mathrm{ff}} = \mathcal{A}_{\mathrm{fm}} = \mathcal{A}_{\mathrm{mm}}, \mathcal{B} = \mathcal{B}_{\mathrm{ff}} = \mathcal{B}_{\mathrm{fm}} = \mathcal{B}_{\mathrm{mm}}$, and $h_{\mathrm{f}} = h_{\mathrm{m}}$, then:

$$F_{\mathrm{IS}} = 1 - \frac{1}{1-x} \tag{S26}$$

and:

$$F_{\mathrm{ST}} = \frac{x}{1 - h^2(1-x)} \tag{S27}$$

Where for the autosomal case:

$$x_A = (1-\mathcal{P})\frac{1}{8}(\mathcal{A}+\mathcal{B}) + \frac{1}{2}\mathcal{P} \tag{S28}$$

For the aphid-like case:

$$x_{X_a} = (1-\mathcal{P})\frac{1}{8}(\mathcal{A}+2\mathcal{B}) + \frac{1}{2}\mathcal{P} \tag{S29}$$

Thus we can see that as $\mathcal{P}$ comes closer to one (meaning that juveniles are likely to descend from the same particular clonal lineage on a patch), then the autosomes and aphid-like X will converge towards the same value for $F_{\mathrm{IS}}$ and $F_{\mathrm{ST}}$. In contrast, when $\mathcal{P} = 0$, i.e. there is no clonal phase, then as $x_A \le x_{X_a}$ then $F_{IS}$ will always be larger (more negative), and $F_{ST}$ will always be larger too, on the aphid-X than on the autosomes.

### 3 Coalescence and low rates of sex

#### 3.1 Constant sex approximation

If sex is very infrequent, then the approximations used above may become less accurate. Instead, gene lineages will take a long time to move between maternal-origin and paternal-origin states, and thus differences will build up between maternal-origin and paternal-origin gene copies, i.e. the Meselson effect. To model this, we now define the matrix $\boldsymbol{T}$ which describes the movement between three types of states: (1) two gene copies residing within an individual, (2) two gene copies residing in different individuals, and (3) coalescence. $\boldsymbol{T}_{\mathbb{s}}$ describes sexual transitions, and $\boldsymbol{T}_{\mathbb{a}}$ describes asexual transitions.

$$\boldsymbol{T}_{\mathbb{s}} = \begin{pmatrix} 1 & 0 & 0 \\ c_{\mathbb{m}}^2\mathcal{A}(1-\mu) + c_{\mathbb{p}}^2\mathcal{B}(1-\nu) & 2c_{\mathbb{m}}c_{\mathbb{p}} + c_{\mathbb{m}}^2(1-\mathcal{A}) + c_{\mathbb{p}}^2(1-\mathcal{B}) & c_{\mathbb{m}}^2\mathcal{A}\mu + c_{\mathbb{p}}^2\mathcal{B}\nu \\ 0 & 0 & 1 \end{pmatrix} \tag{S30}$$

$$\boldsymbol{T}_{\mathbb{a}} = \begin{pmatrix} 1 & 0 & 0 \\ 2c_{\mathbb{m}}c_{\mathbb{p}}\mathcal{P} & 1-\mathcal{P} & (c_{\mathbb{m}}^2 + c_{\mathbb{p}}^2)\mathcal{P} \\ 0 & 0 & 1 \end{pmatrix} \tag{S31}$$

Where:

$$c_{\mathbb{m}} = c_{\mathrm{f}} = \frac{1-\beta}{2-\alpha-\beta} \tag{S32a}$$

$$c_{\mathbb{P}} = c_{\mathrm{m}} = \frac{1-\alpha}{2-\alpha-\beta} \tag{S32b}$$

And $\mathcal{A}$,$\mathcal{B}$, $\mathcal{P}$ are the probabilities of sharing the same parent (female, males, asexual), and $\mu$ and $\nu$ are the probabilities of descending from the same physical gene copy, given that one comes from the same parent. We can then solve for the time to coalescence as before using the approach of Slatkin (Slatkin, 1991; Hartfield et al., 2016; Hartfield, 2021). Let $\boldsymbol{G}$ be the upper left matrix of $\boldsymbol{T}$, which represents the probabilities of non-coalescence.Then our expected coalescence times within and between individuals can be calculated as:

$$\begin{pmatrix} \bar{t}_{\mathrm{w}} \\ \bar{t}_{\mathrm{b}} \end{pmatrix} = (\boldsymbol{I}-\boldsymbol{G})^{-1}\, a(0) \tag{S33}$$

Where:

$$a(0) = \begin{pmatrix} 1 \\ 1 \end{pmatrix} \tag{S34}$$

We can then solve and compare this to the approximation earlier where:

$$\bar{t}_{\mathrm{approx}} = \frac{1}{\bar{Q}} \tag{S35}$$

We can simplify the times to coalescence by expressing them relative to the fast-time scale approximation:

$$\Delta\bar{t}_{\mathrm{w}} = \frac{\bar{t}_{\mathrm{w}} - \bar{t}_{\mathrm{approx}}}{\bar{t}_{\mathrm{approx}}} \tag{S36}$$

$$\Delta\bar{t}_{\mathrm{b}} = \frac{\bar{t}_{\mathrm{b}} - \bar{t}_{\mathrm{approx}}}{\bar{t}_{\mathrm{approx}}} \tag{S37}$$

Which if we assume that there are $N$ individuals in our population, and during the sexual generations there is an even sex ratio, and Poisson distribution of offspring number, then these simplify further. For our standard autosomal systems:

$$\begin{pmatrix} \Delta\bar{t}_{\mathrm{w},A} \\ \Delta\bar{t}_{\mathrm{b},A} \end{pmatrix} = \begin{pmatrix} \frac{1}{\sigma}\left(\frac{1}{4}\sigma(\mathcal{A}+\mathcal{B}) + (1-\sigma)\mathcal{P}\right) \\ \frac{1}{2\sigma}\left(\frac{1}{4}\sigma(\mathcal{A}+\mathcal{B}) + (1-\sigma)\mathcal{P}\right) \end{pmatrix} = \begin{pmatrix} \frac{1}{N\sigma} \\ \frac{1}{2N\sigma} \end{pmatrix} \tag{S38}$$

For our standard X chromosome systems:

$$\begin{pmatrix} \Delta\bar{t}_{\mathrm{w},X} \\ \Delta\bar{t}_{\mathrm{b},X} \end{pmatrix} = \begin{pmatrix} \frac{1}{\sigma}\left(\sigma\frac{1}{9}(4\mathcal{A}+\mathcal{B}) + (1-\sigma)\mathcal{P}\right) \\ \frac{1}{2\sigma}\left(\sigma\frac{4}{9}\mathcal{A} + (1-\sigma)\frac{8}{9}\mathcal{P}\right) \end{pmatrix} = \begin{pmatrix} \frac{9+\sigma}{9N\sigma} \\ \frac{4}{9N\sigma} \end{pmatrix} \tag{S39}$$

And for an aphid like X chromosome:

$$\begin{pmatrix} \Delta\bar{t}_{\mathrm{w},X_a} \\ \Delta\bar{t}_{\mathrm{b},X_a} \end{pmatrix} = \begin{pmatrix} \frac{1}{\sigma}\left(\sigma\frac{1}{4}(\mathcal{A}+\mathcal{B}) + (1-\sigma)\mathcal{P}\right) \\ \frac{1}{2\sigma}\left(\sigma\frac{1}{4}\mathcal{A} + (1-\sigma)\mathcal{P}\right) \end{pmatrix} = \begin{pmatrix} \frac{1}{N\sigma} \\ \frac{2-\sigma}{4N\sigma} \end{pmatrix} \tag{S40}$$

Inspecting these, and visually from Figure S1, we can see that the fast time scale approximation generally works well provided sex is not too infrequent relative to the population size, which for many of the species we describe will be appropriate. Moreover, we can see the convergence between the behaviour between an aphid-like X chromosome and an autosome under high rates of asexuality.

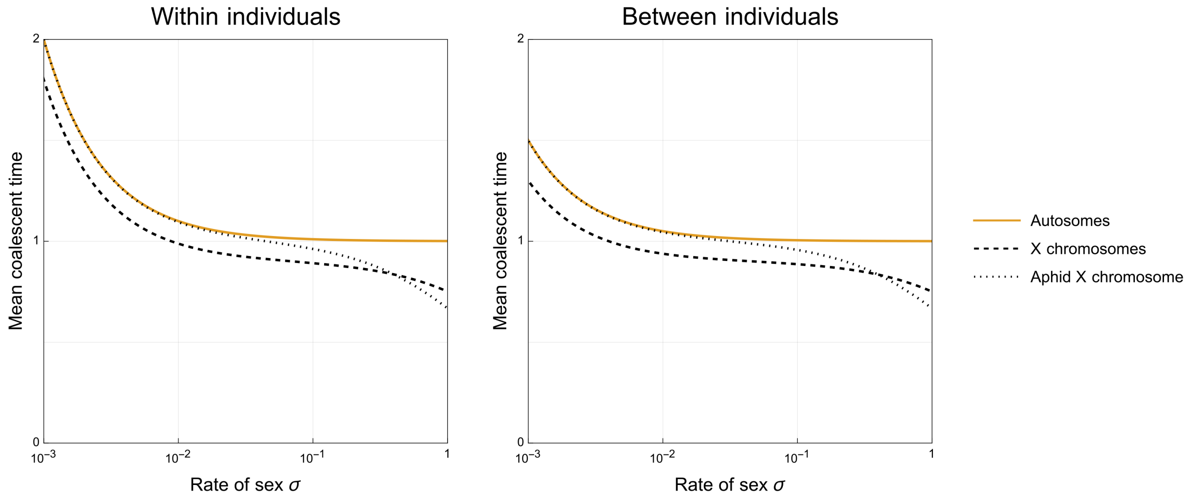


Figure S1: **Mean time to coalescence within and between individuals for different genetic systems and rates of sex.** Coalescent times are expressed in units of $2N$ generations. The population size is $N = 10^3$. During sexual generations an even sex ratio and a Poisson distribution of offspring number is assumed.

#### 3.2 Temporal heterogeneity in rates of sex

Finally, the above analysis approximates the temporal heterogeneity in rates by arithmetically weighting asexual and sexual transitions. We can compare the approximation above, which averages across time, to a numerical solution explicitly considering the temporal structure. Following Hartfield et al. (2016), the probability that two gene lineages coalesce after $t$ generations is:

$$P_{\mathrm{w}}(t) = \begin{pmatrix} 1 \\ 0 \end{pmatrix} \left(\Pi_{t'=0}^{t-1} \mathbf{G}_t\right) \begin{pmatrix} \gamma_{\mathrm{w,t}} \\ \gamma_{\mathrm{b,t}} \end{pmatrix} \tag{S41}$$

$$P_{\mathrm{b}}(t) = \begin{pmatrix} 0 \\ 1 \end{pmatrix} \left(\Pi_{t'=0}^{t-1} \mathbf{G}_t\right) \begin{pmatrix} \gamma_{\mathrm{w,t}} \\ \gamma_{\mathrm{b,t}} \end{pmatrix} \tag{S42}$$

Where $\gamma_{\mathrm{w,t}}$ are the probabilities that two gene lineages at time $t-1$ coalesce at time $t$ given they are within the same individual, or $\gamma_{\mathrm{b,t}}$ in different individuals. These results can be seen in Figures S2 and S3. In general, temporal heterogeneity has a more substantial impact upon the various X chromosomes than on the autosomes.

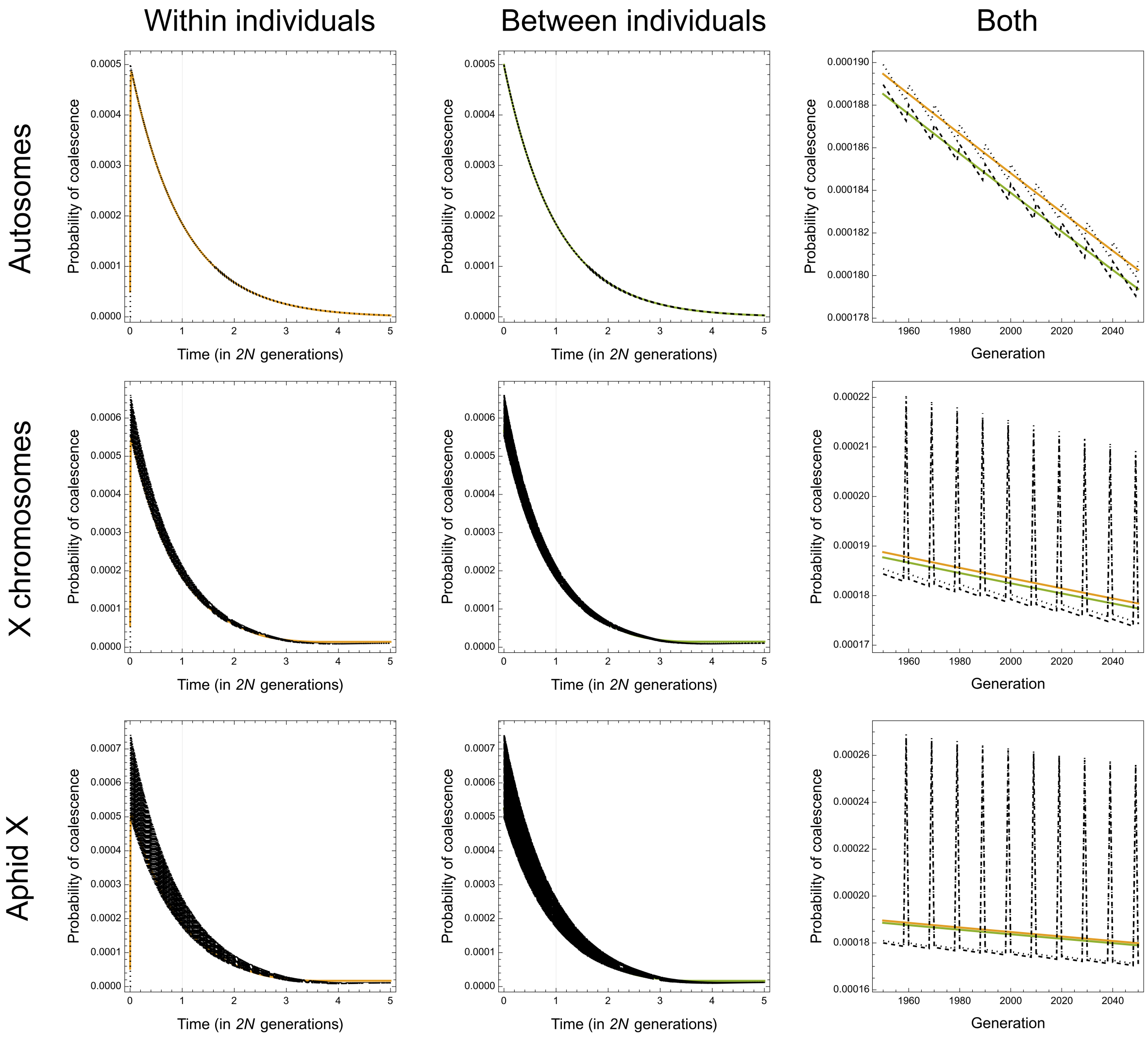


Figure S2: **Probability of coalescence at time $t$ for a population of size $N = 10^3$ and with sex occuring every 10 generations.** Solid lines represent the constant sex approximation, dotted lines include temporal heterogeneity in rates of sex. The rightmost panel is a combination of the two panels to the left, zoomed in around $t = 2000$ to better demonstrate fluctuations in the probability of coalescence caused by temporal heterogeneity in rates of sex.

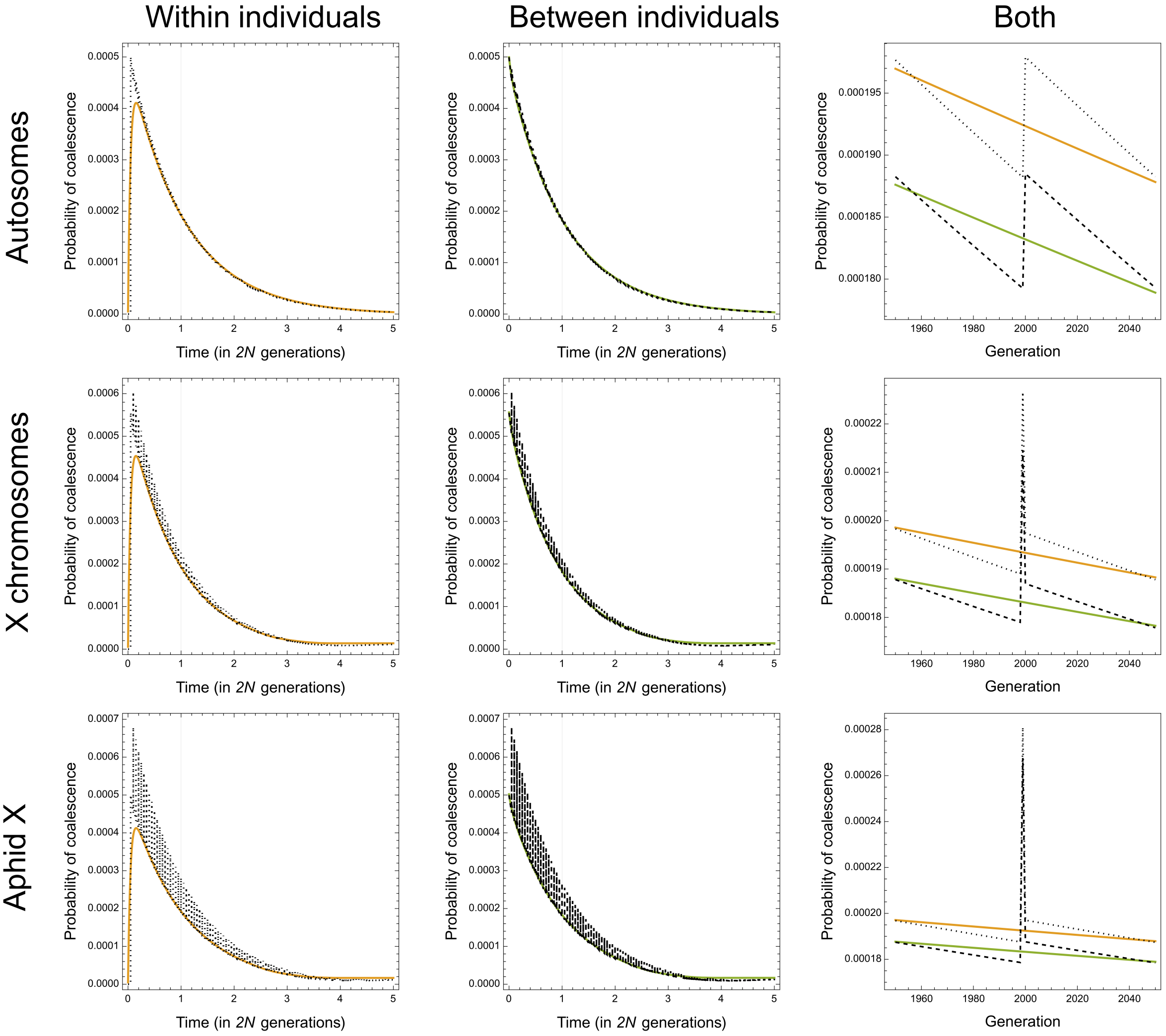


Figure S3: **Probability of coalescence at time $t$ for a population of size $N = 10^3$ and with sex occuring every 100 generations.** Solid lines represent the constant sex approximation, dotted lines include temporal heterogeneity in rates of sex. The rightmost panel is a combination of the two panels to the left, zoomed in around $t = 2000$ to better demonstrate fluctuations in the probability of coalescence caused by temporal heterogeneity in rates of sex.
